## Supporting Information for "The *Salmonella* transmembrane effector SteD hijacks AP1-mediated vesicular trafficking for delivery to antigen-loading MHCII compartments"

Fig S1

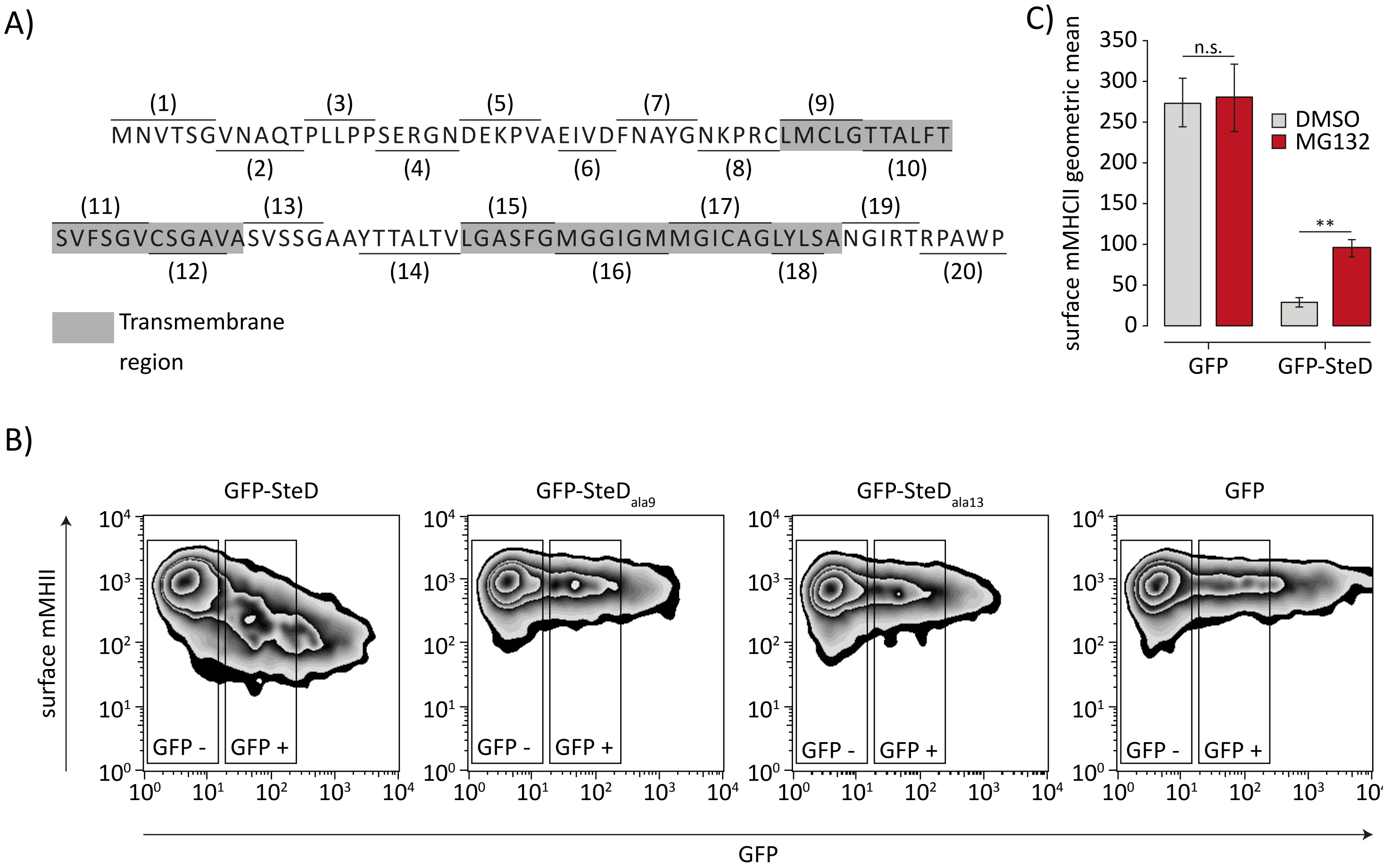


**Fig S1**

(A) Amino acid sequence of SteD showing regions of amino acids substituted to alanine in alanine scanning mutagenesis.

(B) Representative flow cytometry plots showing the gating strategy for GFP-positive cells and negative cells as used for Fig 1C.

(C) mMHCII surface levels of Mel JuSo cells expressing GFP or GFP-SteD and treated with DMSO or MG132 were measured by flow cytometry. Mean of three independent experiments done in duplicate ± SD. Data were analysed by paired t-test, ** $p<0.01$, n.s. – not significant.

Fig S2

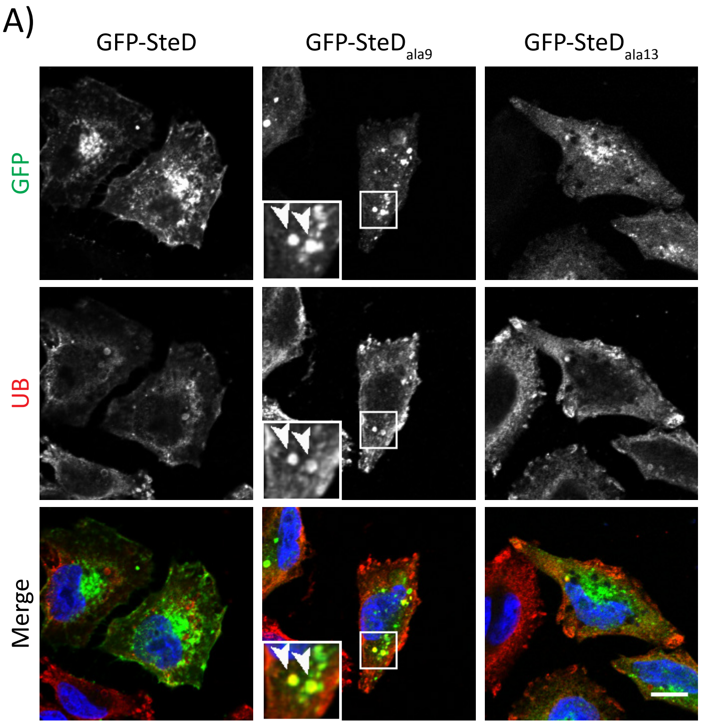


**Fig S2**

(A) Representative confocal immunofluorescence microscopy images of Mel JuSo cells expressing GFP-SteD (wt or mutants) after MG132 treatment. Cells were fixed and processed for immunofluorescence microscopy by labelling for ubiquitin (UB, red), and DNA (DAPI, blue). Arrowheads indicate cellular aggregates. Scale bar – 10 µm.

Fig S3

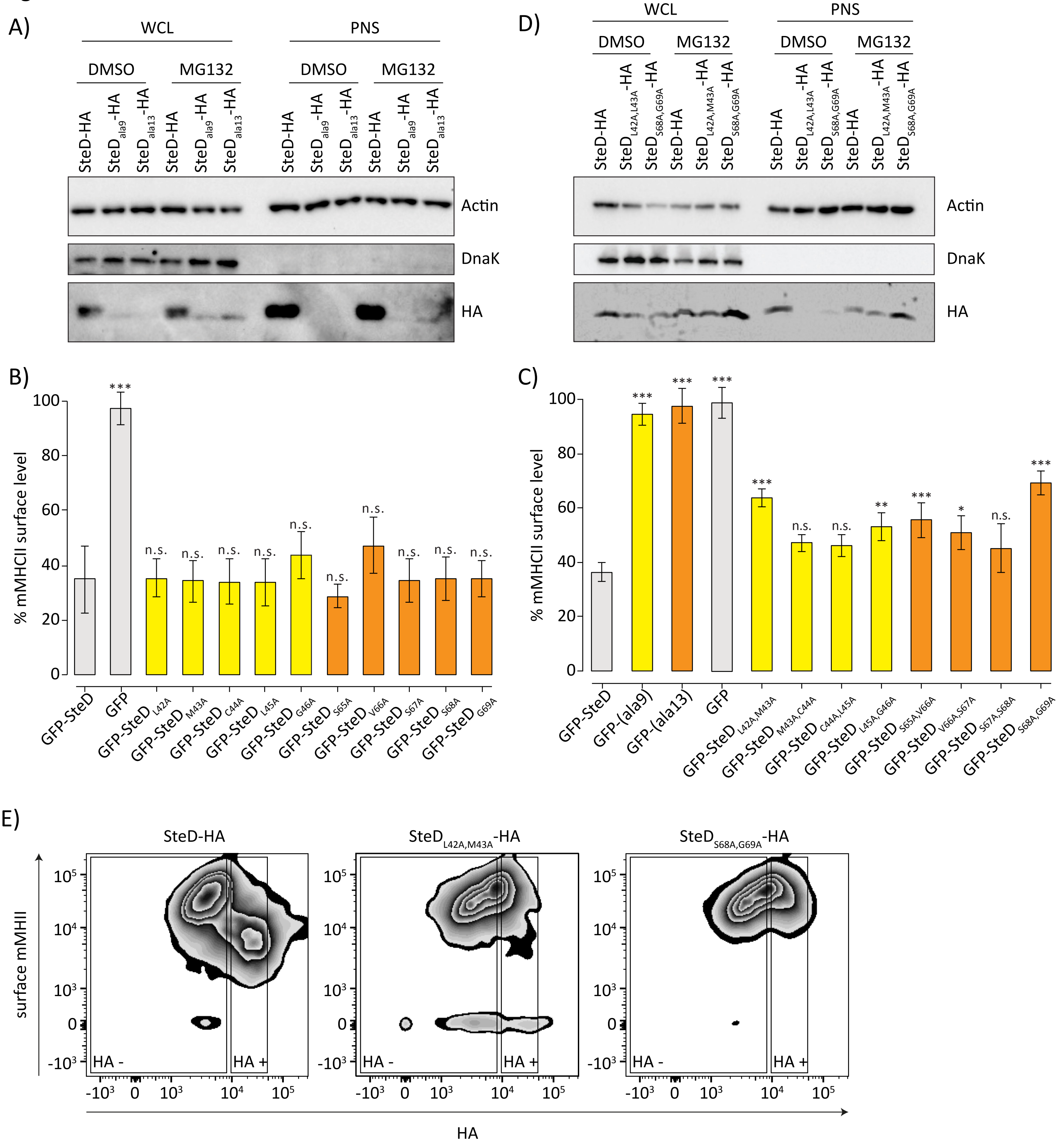


**Fig S3**

(A and D) Protein immunoblots of whole-cell lysates (WCL) and post-nuclear supernatant (PNS) of Mel Juso cells infected with *ΔsteD Salmonella* strains carrying a plasmid expressing SteD-HA (wt or mutants) and treated with MG132 or DMSO carrier. Actin and DnaK represent host cell and *Salmonella* loading controls respectively.

(B and C) mMHCII surface levels of Mel JuSo cells expressing GFP or GFP-SteD (wt or mutants) and treated with MG132. Cells were analysed by flow cytometry and amounts of surface mMHCII in GFP-positive cells are expressed as a percentage of GFP-negative cells in the same sample. Mean of three independent experiments done in duplicate ± SD. Data were analysed by one-way ANOVA followed by Dunnett's multiple comparison test in comparison to wt SteD, *** $p<0.001$, ** $p<0.01$, * $p<0.05$, n.s. – not significant.

(E) Representative flow cytometry plots showing the gating strategy for HA-positive and negative cells as used for Fig 3B.

Fig S4

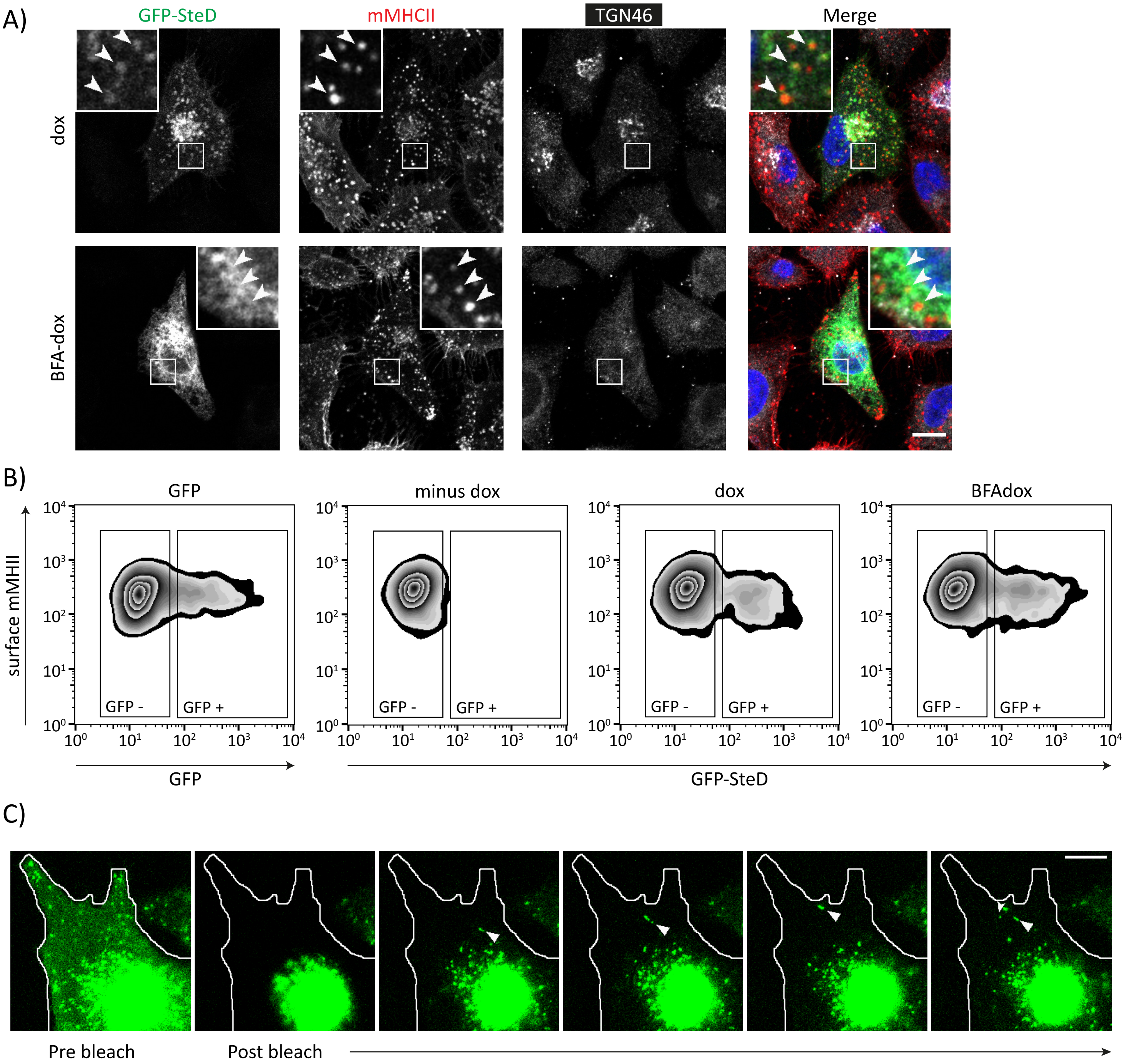


**Fig S4**

(A) Representative confocal immunofluorescence microscopy images of Mel JuSo cells expressing GFP-SteD under a doxycycline-regulated promoter. Cells were either treated with doxycycline for 4 h (dox) or treated with BFA for 3 h followed by doxycycline and BFA for 4 h (BFA-dox). Cells were then fixed and processed for immunofluorescence microscopy by labelling for MHCII compartments (mMHCII, red), the TGN (TGN46, grey), and DNA (DAPI, blue). Arrowheads indicate MHCII compartments. Scale bar – 10 µm.

(B) Representative flow cytometry plots showing the gating strategy for GFP-positive and negative cells as used for Fig 4D.

(C) Confocal microscopy images demonstrating photobleaching of a Mel JuSo cell expressing GFP-SteD from Supplementary Video 2. Closed arrowheads indicate anterograde vesicle traffic from the Golgi region. Barbed arrowhead indicates retrograde vesicle traffic. Scale bar – 10 µm.

Fig S5

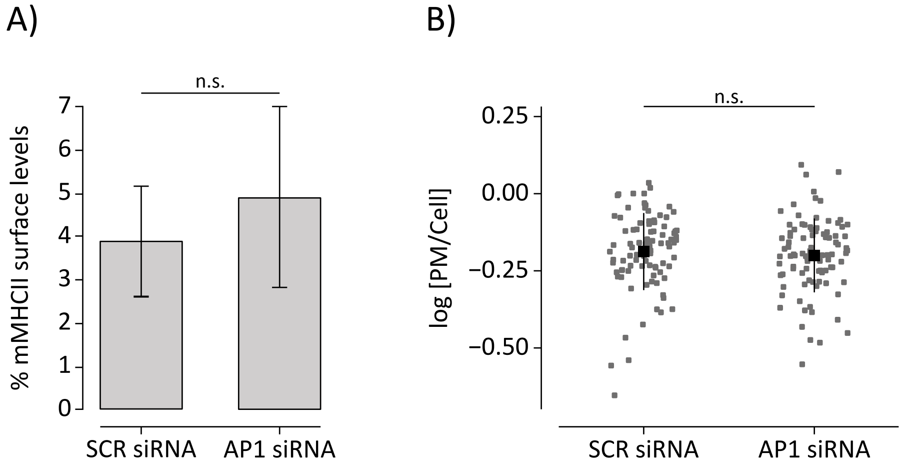


**Fig S5**

(A) mMHCII surface levels of Mel JuSo cells expressing GFP-SteD and treated with scrambled siRNA (SCR) or siRNA specific to the β subunit of AP1. Cells were analysed by flow cytometry and amounts of surface mMHCII in GFP-positive cells are expressed as a percentage of GFP-negative cells in the same sample. Mean of three independent experiments done in duplicate ± SD. Data were analysed by one-way ANOVA followed by Dunnett's multiple comparison test n.s. – not significant. (B) Quantification of GFP at the surface of cells represented in Fig 5D. The fluorescence intensity of the GFP signal at the surface of cells was measured in relation to total cellular fluorescence. Data are representative of three independent experiments. Each dot represents the value for one cell. Mean ± SD. The $\log_{10}$ fold change of the data were analysed by t-test, n.s. – not significant.

Fig S6

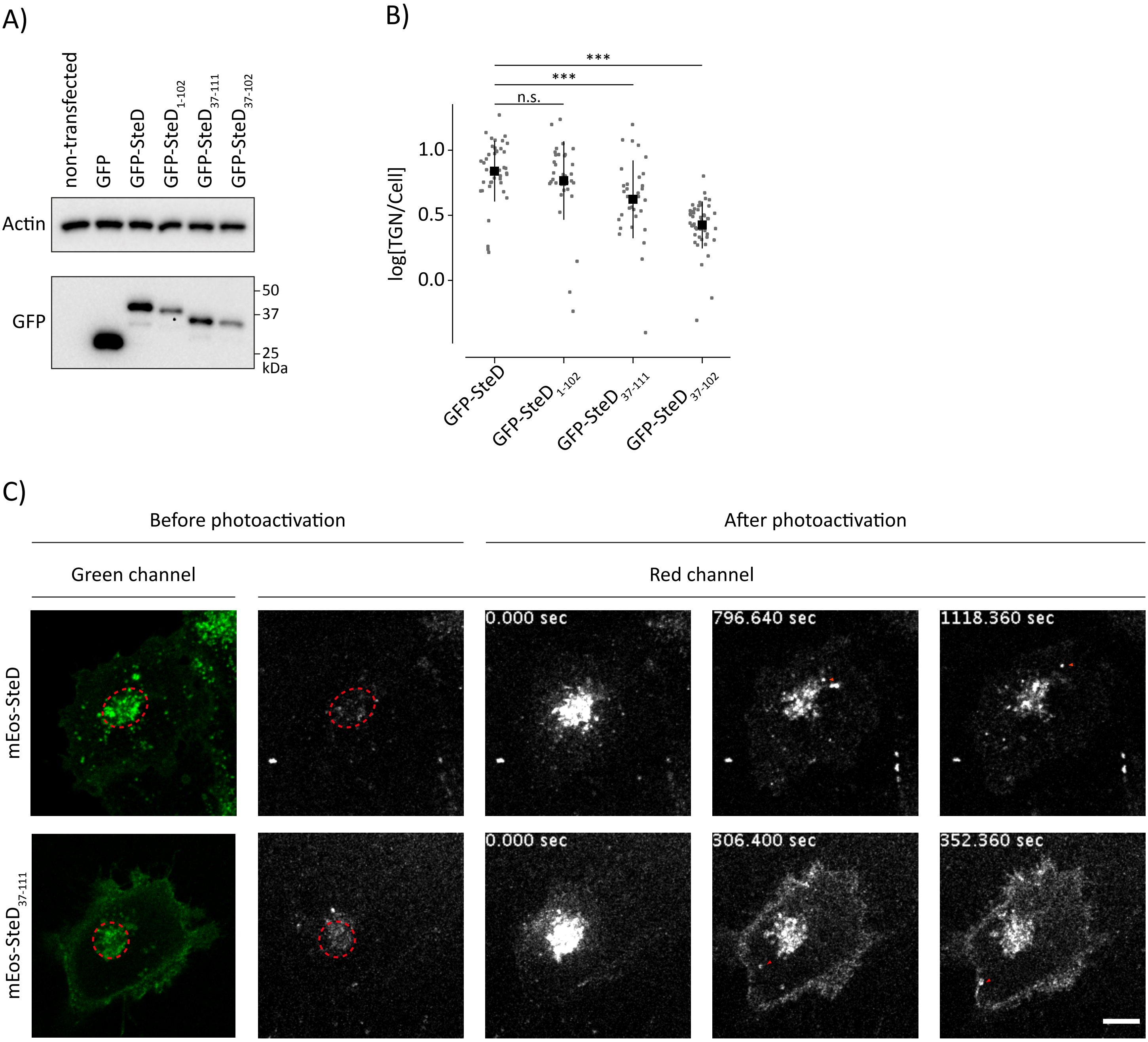


**Fig S6**

(A) Protein immunoblots of Mel JuSo cells expressing GFP or GFP-SteD (wt or mutants).

(B) Quantification of GFP at the TGN of cells represented in Fig 6C. The fluorescence intensity of the GFP signal at the TGN was measured in relation to total cellular fluorescence. Data are representative of three independent experiments. Each dot represents the value for one cell. Mean ± SD. The $\log_{10}$ fold change of the data were analysed by one-way ANOVA followed by Dunnett's multiple comparison test, *** p<0.001, n.s. – not significant.

(C) Confocal microscopy images demonstrating photoactivation of a Mel JuSo cell expressing mEos-SteD (wt or 37-111) from Supplementary Video 3. Red dotted circles indicate photo-activated areas. Red arrowheads indicate Golgi-derived vesicles. Scale bar – 10 µm.

Fig S7

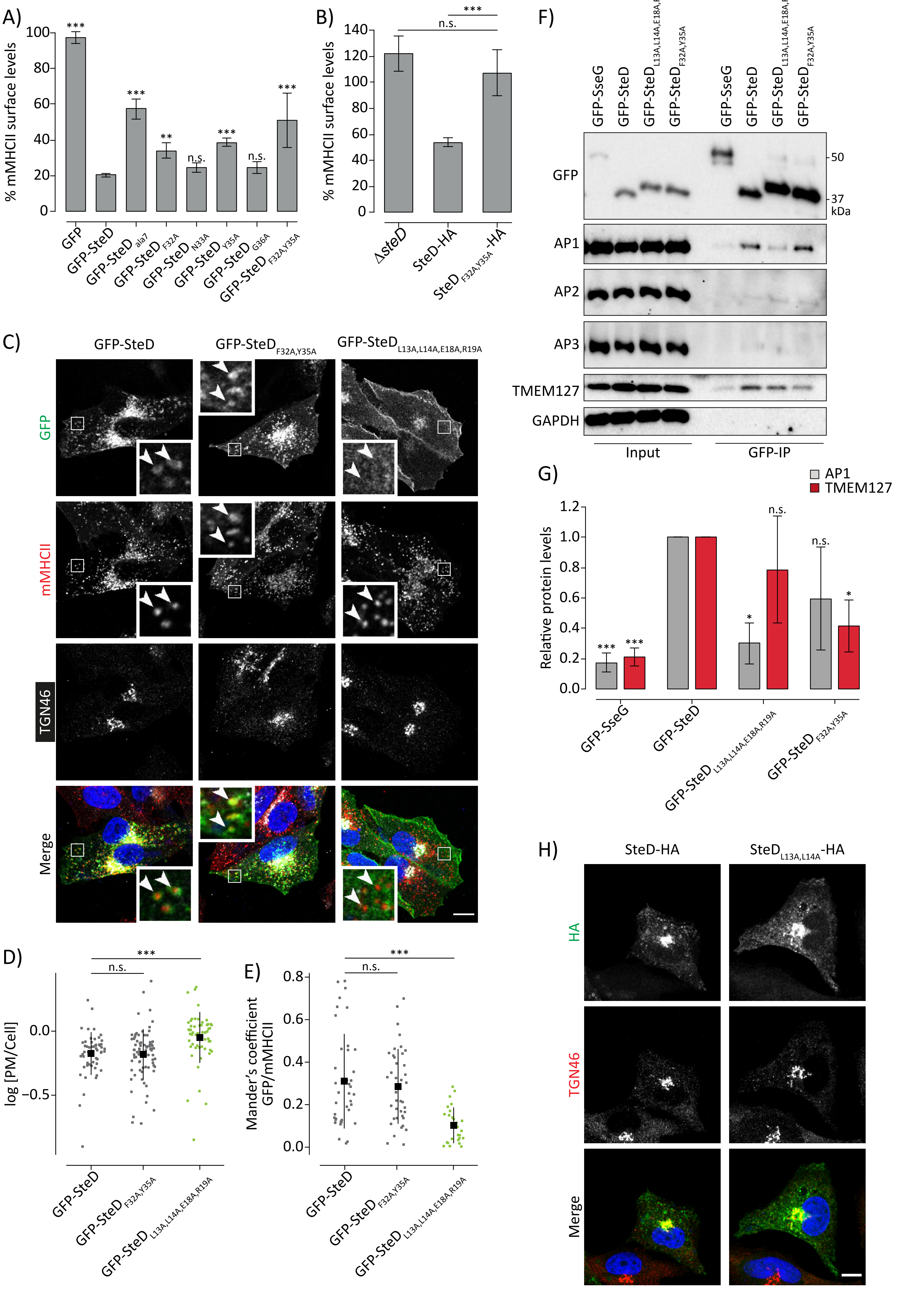
A)
% mMHCII surface levels
100
80
60
40
20
0
***
***
**
n.s.
***
n.s.
***
GFP
GFP-SteD
GFP-SteD ala7
GFP-SteD F32A
GFP-SteD N33A
GFP-SteD Y35A
GFP-SteD G36A
GFP-SteD F32A,Y35A
B)
% mMHCII surface levels
140
120
100
80
60
40
20
0
n.s.
***
ΔsteD
SteD-HA
SteD F32A,Y35A -HA
C)
GFP-SteD
GFP-SteD F32A,Y35A
GFP-SteD L13A,L14A,E18A,R19A
GFP
mMHCII
TGN46
Merge
D)
log [PM/Cell]
0.0
−0.5
n.s.
***
GFP-SteD
GFP-SteD F32A,Y35A
GFP-SteD L13A,L14A,E18A,R19A
E)
Mander's coefficient GFP/mMHCII
0.8
0.6
0.4
0.2
0.0
n.s.
***
GFP-SteD
GFP-SteD F32A,Y35A
GFP-SteD L13A,L14A,E18A,R19A
F)
GFP-SseG
GFP-SteD
GFP-SteD L13A,L14A,E18A,R19A
GFP-SteD F32A,Y35A
GFP-SseG
GFP-SteD
GFP-SteD L13A,L14A,E18A,R19A
GFP-SteD F32A,Y35A
GFP
50
37
kDa
AP1
AP2
AP3
TMEM127
GAPDH
Input
GFP-IP
G)
AP1
TMEM127
Relative protein levels
1.2
1.0
0.8
0.6
0.4
0.2
0.0
***
***
*
n.s.
n.s.
*
GFP-SseG
GFP-SteD
GFP-SteD L13A,L14A,E18A,R19A
GFP-SteD F32A,Y35A
H)
SteD-HA
SteD L13A,L14A -HA
HA
TGN46
Merge

**Fig S7**

(A) mMHCII surface levels of Mel JuSo cells expressing GFP or GFP-SteD (wt or mutants). Cells were analysed by flow cytometry and amounts of surface mMHCII in GFP-positive cells are expressed as a percentage of GFP negative cells in the same sample. Mean of three independent experiments done in duplicate ± SD. Data were analysed by one-way ANOVA followed by Dunnett's multiple comparison test compared to wt SteD, *** p<0.001, ** p<0.01, n.s. – not significant.

(B) mMHCII surface of Mel Juso cells infected with *ΔsteD Salmonella* carrying a plasmid expressing SteD-HA (wt or mutant). Cells were analysed by flow cytometry and amounts of surface mMHCII in infected cells are expressed as a percentage of uninfected cells in the same sample. Mean of three independent experiments done in duplicate ± SD. Data were analysed by one-way ANOVA followed by Dunnett's multiple comparison test, *** p<0.001, n.s. – not significant.

(C) Representative confocal immunofluorescence microscopy images of Mel JuSo cells expressing GFP-SteD (wt or mutants). Cells were fixed and processed for immunofluorescence microscopy by labelling for MHCII compartments (mMHCII, red), the TGN (TGN46, grey), and DNA (DAPI, blue). Arrowheads indicate MHCII compartments. Scale bar – 10 µm.

(D) Quantification of GFP at the surface of cells represented in Fig S7C. The fluorescence intensity of the GFP signal at the cell surface was measured in relation to total cellular fluorescence. Data are representative of three independent experiments. Each dot represents the value for one cell. Mean ± SD. The $\log_{10}$ fold change of the data were analysed by one-way ANOVA followed by Dunnett's multiple comparison test, *** p<0.001, n.s. – not significant.

(E) Mander's overlap coefficient of the fraction of GFP-SteD positive pixels that colocalise with mMHCII positive pixels from cells as represented in Fig S7C. Data are representative of three independent experiments. Each dot represents the value for one cell. Mean ± SD. Data were analysed by one-way ANOVA followed by Dunnett's multiple comparison test, *** p<0.001, n.s. – not significant.

(F) Protein immunoblots of whole-cell lysates (Input) and immunoprecipitation with GFP-trap beads (GFP IP) from Mel Juso cells expressing GFP-SteD (wt or mutants) or GFP-SseG following crosslinking with DSP. Mutation of charged residues might explain the difference in migration through the SDS gel. AP1 – antibody specific for the γ subunit, AP2 – antibody specific for the α subunit, AP3 – antibody specific for the δ subunit.

(G) Levels of immunoprecipitated AP1 and TMEM127 were calculated by densitometry from immunoblots as represented in Fig S7F. Protein levels were normalised to GFP-SteD. Mean of three independent experiments ± SD. The data were analysed by one sample t-test, *** p<0.001, * p<0.05, n.s. – not significant.

(H) Representative confocal immunofluorescence microscopy images of Mel Juso cells infected with *ΔsteD Salmonella* strains carrying a plasmid expressing SteD-HA (wt or mutant). Cells were fixed and processed for immunofluorescence microscopy by labelling for HA (green), the TGN (TGN46, red), and DNA (DAPI, blue). Scale bar – 10 µm.

**Table S1 - *S.* Typhimurium strains used in this study**

| **Name** | **Description** | **Reference** |
|---|---|---|
| wild-type | 14028s *S.* Typhimurium wild-type | ATCC |
| *ΔssaV* | *ΔssaV* ::km | Beuzon et al., 1999 |
| *ΔsteD* | *ΔsteD* ::km | Bayer-Santos et al., 2016 |

Abbreviations: ATCC - American Type Culture Collection, km - kanamycin resistance

**Table S2 - Plasmids used in this study**

| Name | Description | Reference |
|---|---|---|
| m4p GFP | Mammalian retroviral expression plasmid containing GFP | Bayer-Santos et al., 2016 |
| m4p GFP-SteD | Mammalian retroviral expression plasmid containing N-terminal GFP-tagged steD | Bayer-Santos et al., 2016 |
| m4p GFP-SteD$_{ala9}$ | Mammalian retroviral expression plasmid containing N-terminal GFP-tagged steD$_{ala9}$ | Bayer-Santos et al., 2016 |
| m4p GFP-SteD$_{ala13}$ | Mammalian retroviral expression plasmid containing N-terminal GFP-tagged steD$_{ala13}$ | Bayer-Santos et al., 2016 |
| m4p mCherry-SseG | Mammalian retroviral expression plasmid containing N-terminal mCherry-tagged sseG | This study |
| m4p GFP-SteD(SseG TM) | Mammalian retroviral expression plasmid containing N-terminal GFP-tagged chimera of steD with TM domains of sseG | This study |
| m4p GFP-SteD(SseG TM)$_{ala13}$ | Mammalian retroviral expression plasmid containing N-terminal GFP-tagged chimera of steD$_{ala13}$ with TM domains of sseG | This study |
| m4p GFP-SteD$_{LM42AA}$ | Mammalian retroviral expression plasmid containing N-terminal GFP-tagged steD$_{LM42AA}$ | This study |
| m4p GFP-SteD$_{MC43AA}$ | Mammalian retroviral expression plasmid containing N-terminal GFP-tagged steD$_{MC43AA}$ | This study |
| m4p GFP-SteD$_{CL44AA}$ | Mammalian retroviral expression plasmid containing N-terminal GFP-tagged steD$_{CL44AA}$ | This study |
| m4p GFP-SteD$_{LG45AA}$ | Mammalian retroviral expression plasmid containing N-terminal GFP-tagged steD$_{LG45AA}$ | This study |
| m4p GFP-SteD$_{SV65AA}$ | Mammalian retroviral expression plasmid containing N-terminal GFP-tagged steD$_{SV65AA}$ | This study |
| m4p GFP-SteD$_{VS66AA}$ | Mammalian retroviral expression plasmid containing N-terminal GFP-tagged steD$_{VS66AA}$ | This study |

| | | |
|---|---|---|
| m4p GFP-SteD$_{SS67AA}$ | Mammalian retroviral expression plasmid containing N-terminal GFP-tagged steD$_{SS67AA}$ | This study |
| m4p GFP-SteD$_{SG68AA}$ | Mammalian retroviral expression plasmid containing N-terminal GFP-tagged steD$_{SG68AA}$ | This study |
| pWSK29 pSteD-SteD-2HA,SrcA | *Salmonella* expression plasmid containing C-terminal 2HA-tagged steD and srcA including intergenic region with endogenous promoter | Godlee et al., 2019 |
| pWSK29 pSteD-SteD$_{LM42AA}$-2HA,SrcA | *Salmonella* expression plasmid containing C-terminal 2HA-tagged steD$_{LM42AA}$ and srcA including intergenic region with endogenous promoter | This study |
| pWSK29 pSteD-SteD$_{SG68AA}$-2HA,SrcA | *Salmonella* expression plasmid containing C-terminal 2HA-tagged steD$_{SG68AA}$ and srcA including intergenic region with endogenous promoter | This study |
| pcDNA 6/TR | Mammalian expression plasmid for high-level expression of the tetracycline repressor (TR) protein | Life Technologies |
| pcDNA 4/TO GFP-SteD | Mammalian expression plasmid for doxycycline-regulated expression of N-terminal tagged GFP-steD | This study |
| pcDNA 4/TOGFP-SteD$_{ala13}$ | Mammalian expression plasmid for doxycycline-regulated expression of N-terminal tagged GFP-steD$_{ala13}$ | This study |
| m4p GFP-SteD(1-102) | Mammalian retroviral expression plasmid containing N-terminal GFP-tagged steD(1-102) | This study |
| m4p GFP-SteD(37-111) | Mammalian retroviral expression plasmid containing N-terminal GFP-tagged steD(37-111) | This study |
| m4p GFP-SteD(37-111) | Mammalian retroviral expression plasmid containing N-terminal GFP-tagged steD(37-102) | This study |
| m4p mEos3.2-SteD | Mammalian retroviral expression plasmid containing N-terminal mEos3.2-tagged steD | This study with mEos3.2 from Addgene 57484 |
| m4p mEos3.2-SteD(37-111) | Mammalian retroviral expression plasmid containing N-terminal mEos3.2-tagged steD(37-111) | This study with mEos3.2 from Addgene 57484 |
| pWSK29 pSteD-SteD$_{LL13AA}$,SrcA | *Salmonella* expression plasmid containing C-terminal 2HA-tagged steD$_{LL13AA}$ and srcA including intergenic region with endogenous promoter | This study |

| | | |
|---|---|---|
| pWSK29 pSteD-SteD$_{LL/ER}$-2HA,SrcA | *Salmonella* expression plasmid containing C-terminal 2HA-tagged steD$_{LL/ER}$ and srcA including intergenic region with endogenous promoter | This study |
| pWSK29 pSseA-SseF-2HA | *Salmonella* expression plasmid containing C-terminal 2HA-tagged sseF with sseA promoter | Yu et al., 2010 |
| m4p GFP-SteD$_{LL/ER}$ | Mammalian retroviral expression plasmid containing N-terminal GFP-tagged steD$_{LL/ER}$ | This study |
| mCherry-ER-3 | Mammalian expression plasmid for expression of | Gift from Michael Davidson (Addgene 55041) |

**Table S3 - Primary antibodies used in this study**

| Antibody | Source | Use | Dilution |
|---|---|---|---|
| Rat monoclonal anti-GFP clone 3H9 | Chromotek 3H9-100 | WB | 1 in 2000 |
| Rabbit anti-actin | Sigma A2066 | WB | 1 in 2000 |
| Mouse monoclonal HRP-conjugated anti-ubiquitin clone P4D1 | Santa Cruz sc-8017 | WB | 1 in 500 |
| Mouse monoclonal anti-Golgin97 clone CDF4 | eBioscience 14-9767-82 | WB | 1 in 1000 |
| Mouse monoclonal anti-HLA-DR alpha chain clone TAL.1B5 | DAKO M0746 | WB | 1 in 3000 |
| Mouse monoclonal anti-HLA-DR cloneL243 | Sigma-Aldrich | Flow/IF | Flow and IF 1 in 300 |
| Rabbit anti-TGN46 | LSBio LS-B6874 | IF | 1 in 400 |
| Mouse monoclonal anti-ubiquitin clone FK2 | Enzo BML-PW8810 | IF | 1 in 100 |
| Mouse monoclonal anti-DnaK clone 8E2/2 | Enzo ADI-SPA-880-F | WB | 1 in 2000 |
| Mouse monoclonal anti-HA 16B12 | Biolegend 901502 | WB | 1 in 1000 |
| Rat monoclonal anti-HA clone 3F10 | Roche 11867423001 | IF | 1 in 200 |
| Rabbit anti-beta-1 adaptin | Thermo Fisher PA5-66994 | WB | 1 in 500 |
| Mouse anti-gamma adaptin clone 100/3 | Sigma A4200 | WB | 1 in 100 |
| Mouse anti-adaptin alpha clone 8 | BD biosciences 610502 | WB | 1 in 1000 |

| | | | |
|---|---|---|---|
| Mouse anti-adaptin delta clone 18 | BD Biosciences 611329 | WB | 1 in 1000 |
| Rabbit anti-GAPDH | Abcam ab9585 | WB | 1 in 1000 |
| Rabbit anti-TMEM127 | Bethyl Laboratories A303-450A | WB | 1 in 500 |
